# De novo genome assembly and annotation of the saw-scaled viper *Echis carinatus*, a medically important venomous snake in the Indian Subcontinent

**DOI:** 10.1101/2023.09.05.556308

**Authors:** Avni Blotra, Anukrati Sharma, Divya Tej Sowpati, Karthikeyan Vasudevan

**Affiliations:** CSIR- Centre for Cellular and Molecular Biology, Hyderabad, India 500007

**Keywords:** Snakebite, High throughput sequencing, Hybrid assembly, Toxin gene annotation, Venom evolution

## Abstract

*Echis carinatus* is a widely distributed venomous snake in the Indian subcontinent. It is one of the “Big Four” snake species responsible for mortality and severe health complications caused by envenomations. Given the significance of the species for human health, we assembled and annotated the whole genome of *E. carinatus*. Using long reads for assembly and short reads for correction, we achieved a highly contiguous 1.57Gb assembly, with a contig N50 of 17.1 Mb and scaffold N50 of 23.5 Mb. We could map 96.58% of our reads back to the assembled genome, and Benchmarking Universal Single-Copy Orthologs (BUSCO) score of 97.4%, indicates near completeness. 48.59% of the genome had repeat elements. The predominant types of repeats include retrotransposons (LINEs, SINEs and LTR elements), DNA transposons and satellite DNA (SSRs). Additionally, we conducted a gene prediction analysis, identifying a total of 26,711 protein-coding genes. These genes included alternatively spliced products, leading to the prediction of 31,106 proteins, 77.6% of which were successfully annotated using *Blastp* and *InterProScan*. Out of the total predicted gene set, we identified 119 toxin-coding genes, crucial for venom toxin production in the species. The acquisition of toxin sequence-based information resulted in a valuable dataset of genes that are involved in venom production in *E. carinatus*. This dataset advances our understanding of the biological processes underlying venom production and lays the foundation for the development of new antivenom formulations.

## INTRODUCTION

Snakes are limbless reptiles belonging to the suborder Serpentes. Despite undergoing multiple evolutionary changes that led to their widespread distribution, our understanding of snake biology remains poor. There are 3900 recognized, extant snake species in the world, out of which, roughly 600 are venomous (Uetz P. 2022). Snake venom is a mixture of proteins and peptides that act individually or in synergy to incapacitate the prey by acting on a wide range of targets in the body (Xiong & Huang, 2018). Even though all the venomous snakes have the same evolutionary origin, they show a high degree of intra and interspecific venom variation (Chippaux et al., 1991). Extensive homoplasy at the lineage level and rapid adaptations at different geographies make snake venoms an excellent model system to understand evolution of traits under constant natural selection.

Around 10% of the world’s venomous snake species are found in India, and human-snake interactions are inevitable because of the high density of human population. An outcome of this interaction causes envenomation in humans. This poses a significant health challenge across various regions within the Indian subcontinent. A majority of fatal bites are caused by the “Big four”, which are the Indian cobra (*Naja naja*), Russell’s viper (*Daboia russelli*), saw scaled viper (*Echis carinatus*), and common krait (*Bungarus caeruleus*). Because of the knowledge gaps on the venoms and the outcomes of envenomation, snakebites are considered a neglected tropical disease (Waiddyanatha et al., 2019).

*Echis carinatus* has a potent hemotoxic venom that damages tissue and induces symptoms of coagulopathies (Slagboom et al., 2017). Our present understanding of *E. carinatus* venom is only based on venom proteomes, which has provided insight into the complex makeup of venom. Nonetheless, our ability to identify the venom toxins unambiguously is limited, as key isoforms of toxins responsible for diverse pathologies still remain unidentified (Bhatia & Vasudevan, 2020; Casewell et al., 2020). Genomes of venomous snakes are woefully inadequate, thereby a comprehensive library of snake venom toxins has eluded us (Suryamohan et al., 2020).

Therapeutics for snakebite management require a complete understanding of the venom; its composition and the mode of action. Considerable progress has been made in recent years in understanding the composition of snake venoms. Modern whole genome sequencing platforms provide an opportunity to make significant contributions in venom research by providing insights into venom evolution and identification of core toxin coding genes that might improve snakebite treatment strategies.

We used a comprehensive approach involving various genome sequencing platforms and bioinformatic tools to assemble a precise and contiguous de novo whole genome of *Echis carinatus*. The preliminary draft assembly was created using nanopore long read sequences, with subsequent iterative refinement. The errors were removed using long reads followed by polishing with quality-filtered Illumina short read sequences. We also explored the landscape of repeat elements in the genome using repeat masking tools. By implementing a novel assembly strategy, we significantly elevated the accuracy and completeness of the *E. carinatus* genome. Additionally, we conducted RNA-seq analysis of tissue samples and created a well-curated gene library, which was used to predict genes and annotate the genome. This well-annotated genome along with the transcriptome can form the basis for transforming snakebite therapy and genetic studies on snakes.

## MATERIALS AND METHODS

### Sample processing information

Five *E. carinatus* were sourced from different parts of the state of Telangana. Venom glands and other organs were collected after snake’s post-mortem. All snakes were weighed, measured and sexed using protocols described in the MSD Veterinary Manual (https://www.msdvetmanual.com). Genomic DNA was extracted using the Monarch® HMW DNA Extraction Kit for Tissue. To preserve the integrity of RNA, all organs were meticulously collected and stored in RNA later until the RNA extraction process, which was performed using the Qiagen RNeasy kit (Qiagen).

### Genome Sequencing

High molecular weight DNA was extracted from liver of a male *E. carinatus* (EC01) using Monarch® HMW DNA Extraction Kit for Tissue (Cat#T3060L). DNA quantity, purity and integrity were assessed by Qubit fluorometer. Two types of sequencing data were generated: Illumina short-reads and nanopore long reads. For short-reads sequencing, libraries with insert size of 350 bases were prepared and sequenced on an Illumina NovaSeq 6000 platform to generate 150 bp paired-end reads. A total of 710 million short reads were generated. After quality trimming, removing adapters and eliminating the low-quality reads, 107 Gb data remained, which was used for further analysis.

Libraries for long reads were created using a ligation sequencing DNA V14 kit, SQK-LSK-114, using 1μg of high molecular weight genomic DNA as input. The nanopore libraries were sequenced on the PromethION24 sequencer. A total of 94 Gb data, with read N50 of 13.9 kb was generated, giving an overall coverage of 60x based on the estimated genome size. The fast5 files were basecalled into *fastq* format in super accuracy mode using *Guppy v6*.*4*.*6* basecaller. These raw files were quality checked and pre-processed before using it for further analyses.

### RNA-sequencing

Ribonucleic acid (RNA) was extracted using the TRlzol reagent (Invitrogen, USA) followed by the Qiagen RNeasy kit (Cat #74004) from various tissues, including the venom gland, brain, heart, liver, lung, kidney, intestine, testis and ovaries. The quality, purity, and quantity of the extracted RNA were assessed using the HS RNA Qubit fluorometer and the Agilent 4200 tapestation System (Agilent, USA). Subsequently, 1 μg of RNA from each tissue was utilized for the preparation of RNA-sequencing (RNA-seq) libraries with the Illumina stranded total RNA kit. 150 bp paired end reads were generated on the NovaSeq6000 platform (Illumina). For each tissue around 50-60 million reads were generated. Following the quality check and control of the RNA-seq data, we employed the Hisat2 tool to map the data on the genome.

### Quality check of raw reads

#### DNA Seq

To construct a high-fidelity genome, we employed both long and short-read third-generation sequencing technologies. An initial assessment revealed that ONT reads had an average q score of 15.3, with 97.6% of them exceeding the default quality threshold of 12. We utilized *Porechop v0*.*2*.*4* (Wick et al., 2017) and *Nanofit v2*.*8*.*0* (De Coster & Rademakers, 2023) to remove adapters and filter high quality reads, with a cutoff set at 12. These preprocessing steps boosted the quality to 17.3, enhancing the reliability of subsequent analyses. For short reads, *Cutadapt* v4.0 (Martin, n.d.) was employed to eliminate poly A’s and G’s, which were overrepresented in the *fastQC* report. *MultiQC* report reads with quality scores <25 was removed using a *Cutadapt* quality score filter along with the adapter sequences.

#### RNA Seq

From five *E. carinatus* - heart, kidney, brain, venom gland, intestine, lungs, liver, testis, and ovaries were collected. RNA from the tissues were isolated and sequenced. A quality assessment showed the total reads ranged from 40 to 60 million across samples. *Cutadapt* was applied to filter reads, eliminating adapters and employing a quality score cutoff of 25. Hisat2 was utilized to align RNA sequences from these tissues to the genome, with consistent mapping percentages exceeding 80% for each sample.

### Estimation of genome size and coverage

The genome size was estimated using a K-mer frequency-based method. Using *Kmergenie* (Marçais & Kingsford, 2011) we arrived at the best K-mer size and counted the unique kmers using the *Jellyfish* 2.3.0. *GenomeScope* was used to determine the frequency of K-mers, the genome size and the coverage.

### De novo genome assembly

To assemble the genome of *E. carinatus, Flye 2*.*9*.*2* (Kolmogorov et al., 2019) was used with regular ONT data “--nano-raw” that works well for the FLO-PRO114M, R10.4.1 chemistry flowcells. The assembly was enhanced by the “-scaffold” parameter, a feature within *Flye*, which introduced Ns (nucleotide placeholders) to arrange contigs into coherent scaffolds. A statistical summary of assembly quality was generated using *QUAST 5*.*2*.*0* (Gurevich et al., 2013).

### Statistics to assess assembly quality

Numerical statistics, such as N50 are often misleading and could lead to erroneous assessment of the quality of assembly. Therefore, we used a robust approach using biological information-based comparison of multiple assemblies, widely referred to as Benchmarking Universal Single-Copy Orthologs (BUSCO). We used this alignment against the vertebrate lineage database to look at the coverage of the orthologs in our assembly. Furthermore, the accuracy of the assembly and the overall quality was quantified using *Inspector v1*.*0*.*2* (Chen et al., 2021). It identified errors after mapping of the long reads over the contigs. Subsequently, a polishing procedure using *Pilon* v1.24 (Walker et al., 2014) with high quality illumina data was carried out.

### Repeat element identification

The simple repeat sequences (SSRs) and all tandem repeat elements in the whole genome were annotated using *RepeatModeler v2*.*0*.*4* (https://github.com/Dfam-consortium/RepeatModeler). The *de novo* predictions were made using a custom Dfam library -an open database of transposable elements families, *TRF*-tandem repeat finder and *RepeatScout* 1.0.6 - De novo repeat finder. The constructed library was given as an input for *RepeatMasker v4*.*0*.*7* (http://www.repeatmasker.org) to predict the repeats, which were masked using a combination of de novo and annotated repeat databases for downstream analyses

### Gene prediction and annotation

We used the *BRAKER v3*.*0*.*7* annotation pipeline for comprehensive gene annotation. It employs GeneMark-ETP and AUGUSTUS for accurate gene training and prediction and integrates de novo prediction, homology-based searches, and transcriptome/proteome-assisted methods. Additionally, a custom database subset of Sauropsida in the Vertebrata database was incorporated to enhance annotation efficiency. For a comprehensive functional annotation of genes, we employed the *Blastp* function using *Diamond v2*.*0*.*9*.*147*. Hits were then filtered using the following parameters, E-value threshold of 1.0 x 10^-4^, a query coverage > 70% and identity >70%. We then aligned the protein sequences from the finalized gene set to the GenBank *Non-Redundant Protein Database (NRDB)* to reveal sequence similarities and potential functional associations. We incorporated *InterProScan* to further enrich our annotation. This application aided in predicting essential motifs and domains, and facilitated the retrieval of Gene Ontology (GO) terms, to create a nuanced and informative characterization of the genes. To investigate the presence of potential toxin-coding genes within our predicted set, we utilized the Animal Toxin Annotation Project (Tox-Prot) database. We applied two distinct filtering criteria: Relaxed criteria, with query coverage >50%, identity > 50% and an E-value <1 x 10^-4^, and more stringent criteria, identity > 80% and query coverage >80%.

## RESULTS

### De novo genome assembly

The genome size of *E. carinatus* was estimated to be 1.57Gb with an optimal K-mer size of 99 as reported by *Kmergenie*. The coverage was calculated to be 60x, as the available long read data size was approximately 94Gb. To generate a contiguous assembly, we filtered out the small and fragmented sequences. We used the reads that were greater than 10 Kb in length and it resulted in a reduction of sequence coverage to 33x. However, this enhanced the N50 metric. The final assembly was 1.57 Gb in size with 910 contigs. After refinement the assembly had 711 contigs that improved the N50 from 17.1 Mb to 23.5 Mb. 90% of the assembly had 70 contigs (L90) and 99.3 % of the assembly contained contigs over 50kb.

### Assembly Evaluation and correction

The assembly completeness score (BUSCO) was 97.4%. Out of the total 3,327 conserved genes screened in the vertebrate lineage database, 3,239 were complete, 25 were fragmented, and 62 were missing BUSCOs (Table 1). *Inspector* tool was used to map long reads onto assembly contigs to identify errors. This step achieved a mapping rate of 96.58% and initially reported a Quality Value (QV) of 33.2. Additionally, it detected 254 minor assembly errors per million base pairs and introduced 113 Ns during scaffold-level assembly. The approach of correcting the assembly using long reads enhanced the QV to 36.7 and reduced the number of Ns to 33. The minor assembly errors were brought down to 36 per Mbp. After further polishing using the short illumina reads, the assembly metrics had 711 contigs with N50 of 23.5 Mb, and L90 of 69.

**Table 1:** Representing the BUSCO completeness statistics of the *Echis carinatus* genome assembly using vertebrate lineage database.

| **BUSCO Statistics** |  |
| --- | --- |
| Complete BUSCO | 3267 (97.4%) |
| Complete and single copy BUSCOs | 3239 (96.6%) |
| Complete and duplicated BUSCOs | 28 (0.8%) |
| Fragmented BUSCOs | 25 (0.7%) |
| Missing BUSCOs | 62 (1.9%) |
| Total BUSCO groups searched | 3354 |

### Repeat Annotation

We were able to mask 6.74% of repeats in the genome with the Viperidae repeat database. We could construct a custom library with Dfam and Viperidae database and mask 48.59% of repeats (Table 2; Fig. 3). 3.4 % SSRs were masked using our in-house tool DiviSSR v0.1.2 (Avvaru et al., 2021).

**Table 2:** Repeat element distribution in *Echis carinatus* genome.

| **Repeat element distribution** |  |
| --- | --- |
| **Retrotransposons** | **21.88%** |
| LINEs | 15.9% |
| L2/CR1/Rex | 6.9% |
| R2/R4/NeSL | 0.66% |
| RTE/Bov-B | 4.1% |
| L1/CIN4 | 2.8% |
| LTR elements | 5.98% |
| BEL/Pao | 0.06% |
| Ty1/Copia | 0.65% |
| Gypsy/DIRS1 | 4.98% |
| Retroviral | 0.29% |
| **DNA transposons** | **0.79%** |
| hobo-Activator | 0.13% |
| Tc1-IS630-Pogo | 0.65% |
| **Unclassified** | **23.63%** |
| **Satellites** | **2%** |

**Fig. 1:**
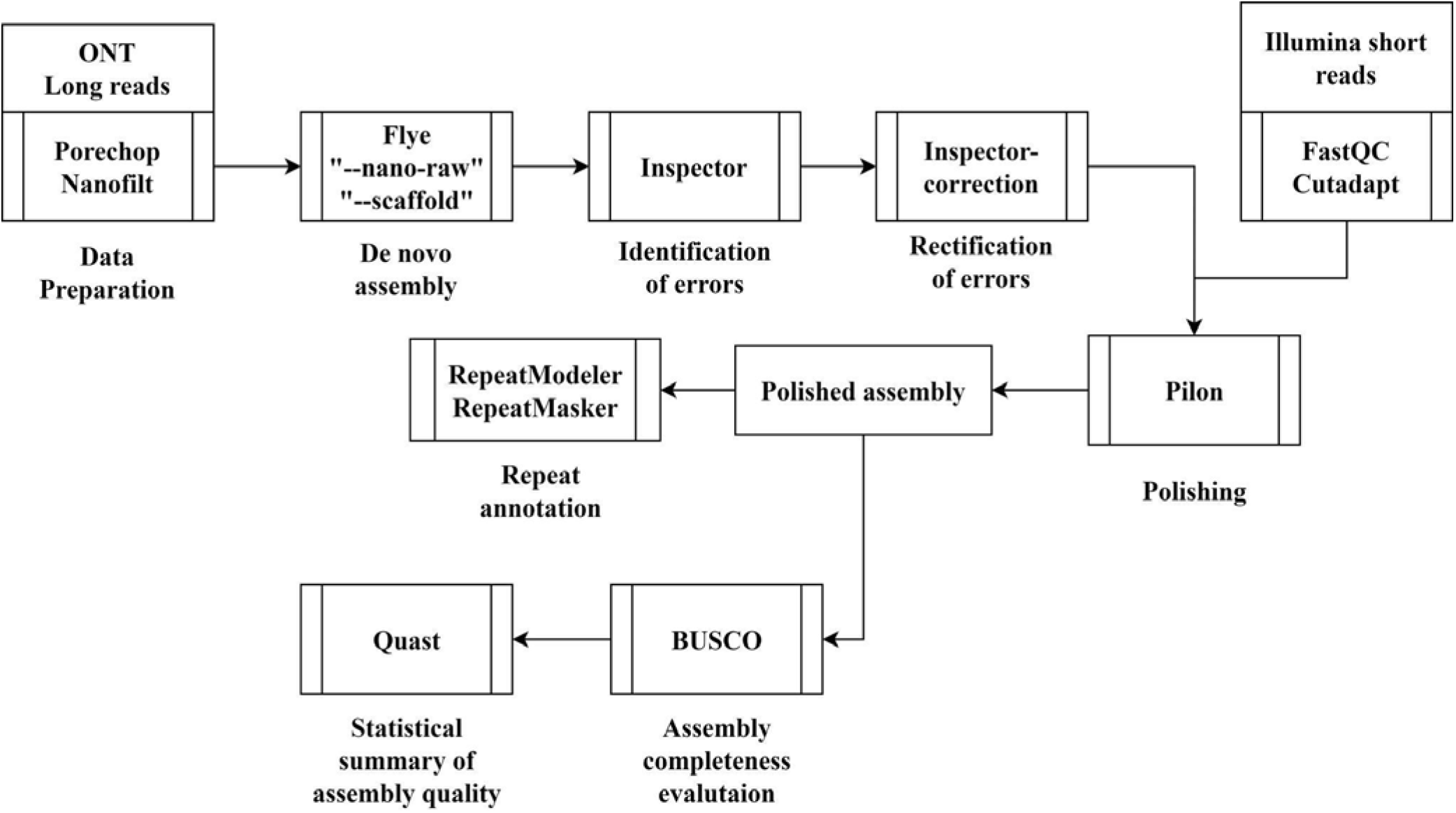
Overview of the de novo assembly pipeline representing the steps and the tools involved.

**Fig. 2:**
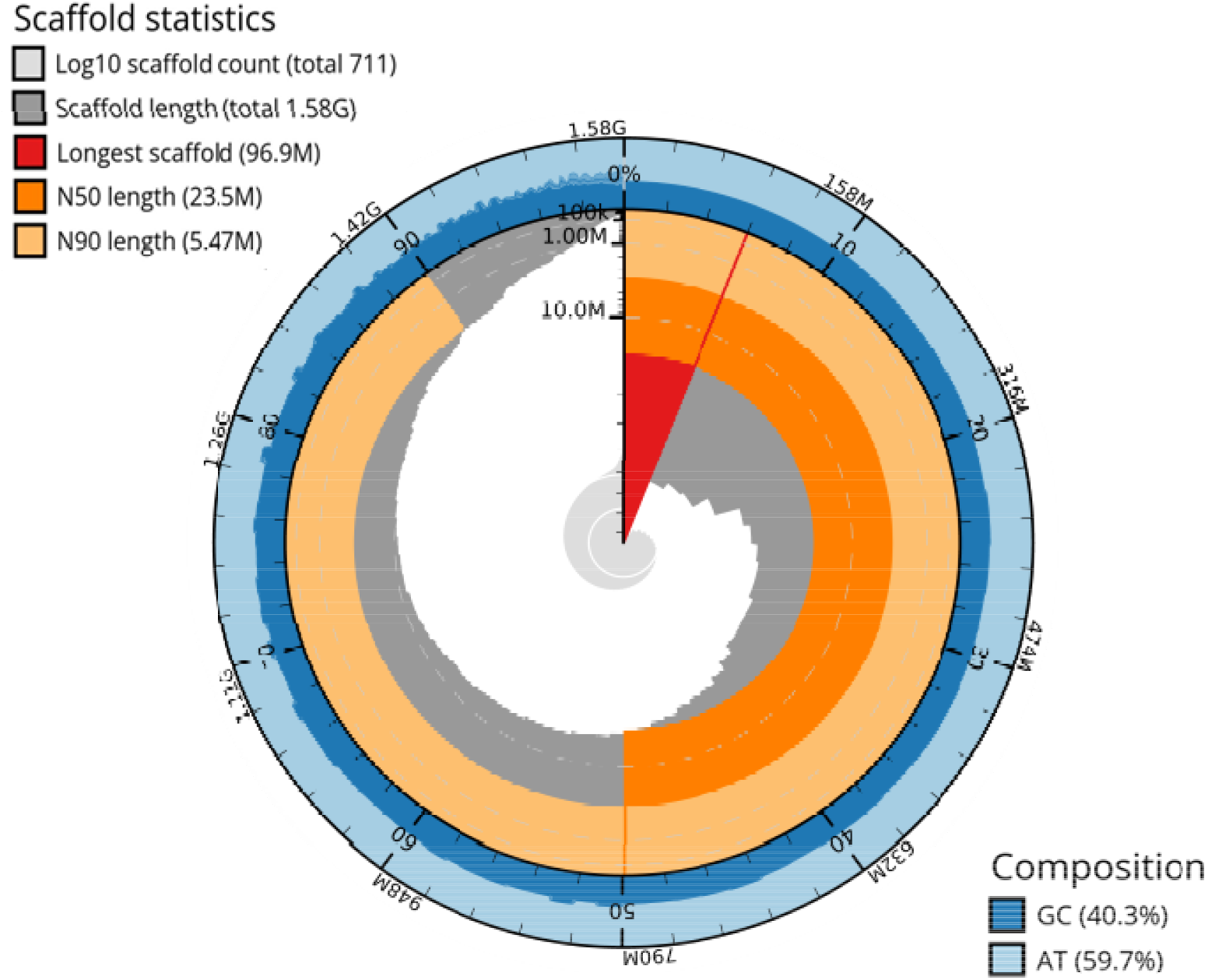
Snail plot summarizing metrics of the *E. carinatus* genome including the length of the longest contig (96.9 Mb; red line), N50 (23.5 Mb; dark orange), N90 (5.47 Mb; light orange), and base composition.

**Fig. 3:**
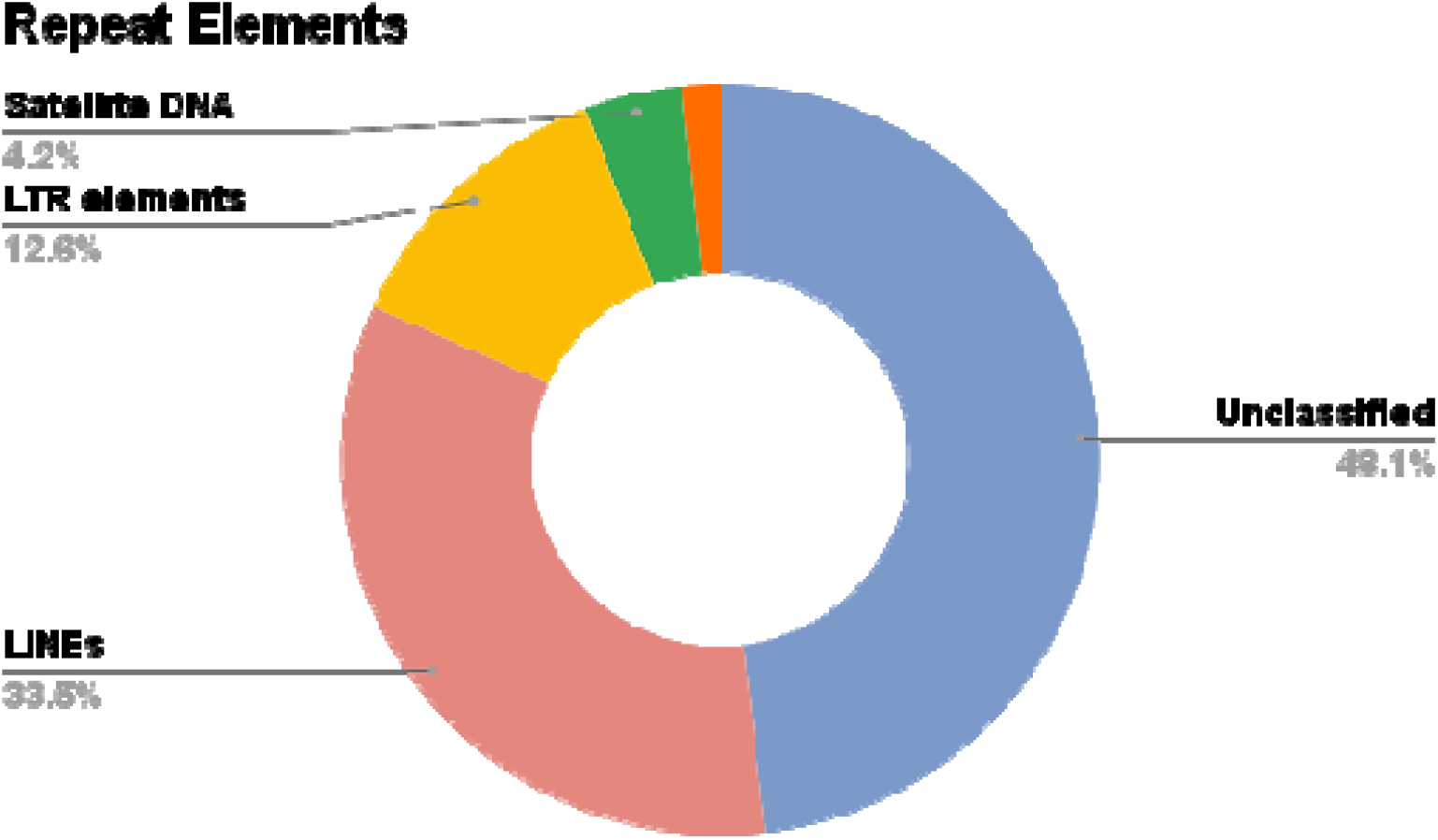
Plot displaying the predominant types of repeat elements present in *Echis carinatus* genome.

### Gene prediction and annotation

We used *Braker* pipeline to annotate the genome with gene expression data from 8 different tissues. We predicted 26,711 protein-coding genes, with 74.8% supported by either RNA-Seq data or protein databases. This gene set encoded a total of 31,106 predicted proteins including the alternatively spliced products, each with corresponding orthologs in NCBI’s non-redundant database. *Blastp* analysis facilitated the successful annotation of 22,219 transcripts (71.4%), corresponding to 20,113 distinct genes. Subsequently, integrating *InterProScan* resulted in the successful annotation of 24,159 transcripts that improved the annotation from 71.4% to 77.6%. Notably, 29,331 transcripts (94.2%) had canonical start and stop codons.

The genome assembly consisted of 711 fragments, out of which, 678 were contigs and 33 were scaffolds. We could map 54.6 % of functionally annotated genes on 32 scaffolds and the remaining genes on 186 contigs. We initially identified 4,407 toxin coding transcripts that met the relaxed criteria out of the 7,839 protein sequences in Tox-Prot. By employing a stringent criteria, we identified 119 toxin-coding genes, crucial for venom toxin production in the species. The completeness of the annotation was 97.6% with the Vertebrate database and 95% with the Sauropsida database (Table 3).

**Table 3:** Representing the BUSCO completeness statistics of the *Echis carinatus* transcriptome using two lineage databases.

|  | **Vertebrate database** | **Sauropsida database** |
| --- | --- | --- |
| Complete BUSCO | 3274 (97.6%) | 7105 (95%) |
| Complete and single copy BUSCOs | 2877 (85.8%) | 5907 (79%) |
| Complete and duplicated BUSCOs | 397 (11.8%) | 1198 (16%) |
| Fragmented BUSCOs | 25 (0.7%) | 42 (0.6%) |
| Missing BUSCOs | 55 (1.7%) | 42 (4.4%) |
| Total BUSCO groups searched | 3354 | 7480 |

### Future Perspective

The Indian subcontinent faces a huge burden of snakebite deaths (Suraweera et al., 2020). Out of the common four venomous snake species, only *Naja naja* has whole genome and transcriptome data. The lack of genomic and transcriptomic data has impeded our understanding of venom toxin evolution and the mechanisms that regulate toxin-gene expression. With the *E. carinatus* genome and transcriptome, we will perform differential expression analysis and refine the venom proteome using a species-specific protein database. This would provide insights into the molecular basis of venom production. More importantly, an in-depth understanding of the toxin gene sequences will lay the foundation for developing targeted therapies for snake bite envenomations.

## Data availability

Raw and assembled sequence data will be available via the NCBI GEO

## Acknowledgements

We are grateful to the Telangana Forest Department for permitting us to collect saw scaled viper and its venom under order no – Rc. No.26803/2012/WL-2 dated 08.04.2021. A. Srinivas and N. Tulasi from the CCMB’s NGS facility generated the sequencing data. Sampath Kumar, Alka Sahu, Gopi Krishnan, Omkar Yadav, K. Rajyalakshmi and Harika Katakam extended help for this work. Anita Malhotra facilitated AB’s research through Bangor University, Taith Research Mobility Scheme for student exchange grant. Aaron Comeault and Axel Barlow provided bioinformatics training for the whole genome analysis.

